# A Hierarchical Model of Purinergic Receptor Activation in Bronchial Epithelial Cells

**DOI:** 10.64898/2026.06.20.733544

**Authors:** Volker Meidl, Martina Kiefmann, Torsten Goldmann, Christian Börnchen, Rainer Kiefmann

## Abstract

Purinergic signaling coordinates diverse epithelial responses to extracellular nucleotides such as ATP, ADP, and UDP. Although many epithelial cell types co-express multiple P2 receptors, the logic by which these receptors integrate nucleotide signals has remained unclear. Here, using primary human airway epithelial cells as a model, we reveal a hierarchical system in which P2Y_2_ functions as a central licensing receptor that both enables and constrains downstream activation of P2Y_6_ and P2Y_12_. Molecular analysis, calcium assays, and pharmacological profiling show that P2Y_6_ and P2Y_12_ exhibit intrinsic activity when co-express to P2Y_2_ but in turn lose responsiveness to their specific agonists upon upstream activation of P2Y_2_. This gating mechanism filters background noise by secondary nucleotides and enforces contextual control over downstream signaling. These findings uncover a previously unrecognized principle of purinergic receptor coordination that may apply broadly across epithelial systems, and offer new insight into nucleotide signaling as a therapeutic target.

## Introduction

Purinergic signaling is a fundamental regulatory mechanism in epithelial biology across diverse tissues [1, 2]. Particularly the P2Y family is of importance in epithelial cells, where it mediates various physiological processes, including ion transport [3], mucus secretion [4, 5] ciliary beat regulation [6], and inflammatory signaling [7]. The P2Y receptor family comprises two subgroups—P2Y1-like (P2Y_1,2,4,6,11_) and P2Y12-like (P2Y_12,13,14_)—characterized by high sequence homology [8]. These G protein–coupled receptors are activated by extracellular nucleotides (ATP, UTP, ADP, UDP, nucleotide sugars) and couple predominantly to Gq/11, Gs, or Gi proteins at the intracellular plasma membrane [9, 10].

Purinergic signaling in epithelia cells is viewed as a tiered system. Nucleoside triphosphates (NTPs), such as ATP, are released in response to mechanical stress, cell injury, or infection [11–14], and are sequentially degraded into nucleoside diphosphates (NDPs), monophosphates, and ultimately nucleosides [15]. These metabolites activate distinct sets of receptors, resulting in a temporally and spatially regulated signaling cascade. For example, ATP activates P2Y_2_, P2Y_4_ and P2Y_11_ (and certain P2X receptors); ADP activates P2Y_1_, P2Y_12_ and P2Y_13_ [16]; and adenosine signals through P1 receptors [17]. In addition, NDPs or adenosine can be released independently, primarily originating from intracellular stores [18].

However, such a model assumes that all purinergic receptors may operate independently, simply responding to the presence or absence of their respective ligands. In contrast, we hypothesize that purinergic receptors constantly operate within a hierarchical regulatory framework, in which the activation or signaling status of one receptor (e.g., P2Y_2_) conditions the responsiveness of others (e.g., P2Y_1,12_). This would allow the cell to execute conditional logic, adding a layer of biological control that prevents inappropriate downstream activation by secondary nucleotides in the absence of a primary receptor target signal.

P2Y_2_ is widely recognized as the principal purinergic receptor in airway epithelial physiology, driving robust calcium signaling in response to ATP and UTP [19, 20]. Limited genetic and pharmacological studies suggest that P2Y₂ accounts for the majority of all purinergic calcium responses in this tissue [21], despite the co-expression of other P2 receptor subtypes [22], raising important questions about how their functions are integrated within the same cellular context.

To investigate this, we studied the functional relationship between P2Y_2_, P2Y_6_, and P2Y_12_ that were intrinsically active in our human primary bronchial epithelial cells (hBECs). Cells were isolated from residual donor lung tissue, as well as obtained from commercial primary cultures. Both submerged and air-liquid interface (ALI) culture models were used to simulate physiological and differentiated epithelial states. This cell type expresses a wide array of purinergic receptors and represents a relevant model to study purinergic coordination in the airway.

Our results reveal that P2Y_2_ activation by ATP/UTP initiates a robust Ca²⁺ signal as previously reported. Unexpectedly, we find that pre-stimulation, inhibition, or knockdown of P2Y_2_ abolishes functional responses mediated by P2Y_6_ and P2Y_12_, despite the presence of their respective ligands. Our findings reveal that P2Y₂ not only serves as the dominant receptor for ATP/UTP-induced calcium responses but also plays a regulatory role as a licensing receptor that dictates the activity of P2Y₆ and P2Y₁₂. These data provide new insights into the hierarchical organization of purinergic signaling in the airway epithelium, and support the idea that purinergic signaling operates not as a set of parallel pathways, but through a hierarchical logic that fine-tunes epithelial responses to extracellular nucleotides.

## Results

### Expression of Purinergic Receptors in Primary Human Bronchial Epithelial Cells

We first screened for P2 receptor (P2R) expression in hBECs cultured under submerged conditions. At the mRNA level, we detected transcripts for P2X_5_ and P2X_6_, as well as for P2Y_2_, P2Y_6_, P2Y_12_, and P2Y_14_ (Fig. 1a). Protein expression largely confirmed the transcript data, with detectable levels of P2X_1_, P2X_4_, P2X_5_, and P2X_6_, and P2Y_2_, P2Y_6_, P2Y_12_, and P2Y14 (Fig. 1b). The specificity of the antibody signals was validated using either blocking peptides or IgG controls (Fig. 1c). These results established the presence of multiple ATP-, UTP-, ADP, and UDP-responsive P2Rs in hBECs.

**Fig. 1.**
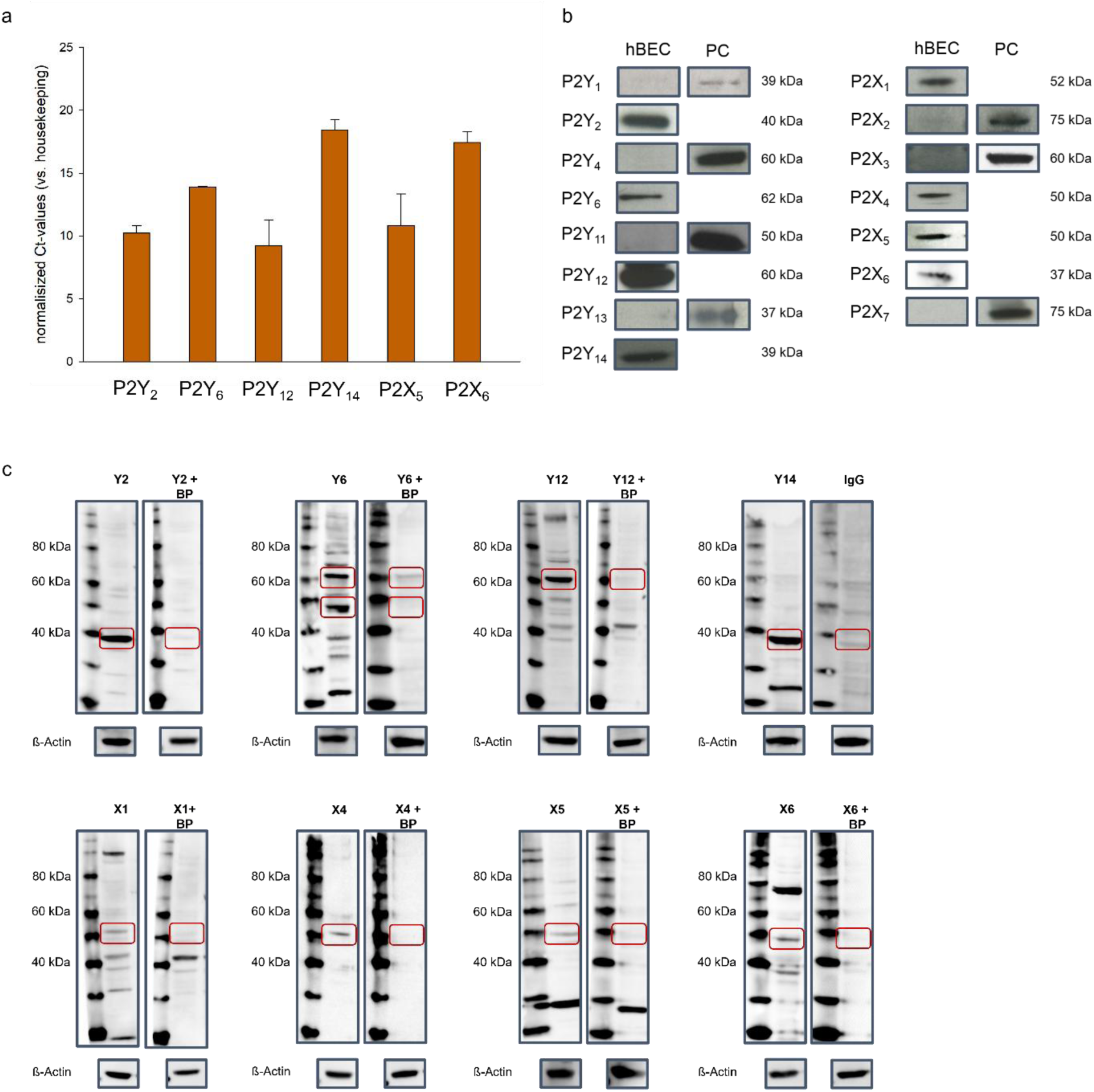
**a** Expression of P2Y and P2X receptors at the mRNA level in hBECs assessed by RT-qPCR; RPL13A-1 served as housekeeping gene (n = 3 experiments per receptor). **b** Representative western blots of P2Y and P2X receptors in hBECs or in positive control cell lines (RLE-6TN: P2Y_1_, P2Y_13_, P2X_3_; R3-1: P2Y_4_; L2: P2Y_11_; C38: P2X_2_; pAE: P2X_7_); black bands indicate receptor-specific antibody binding (n = 3 experiments per receptor). **c** Representative western blots of P2Y and P2X receptors from panel b, including blocking peptides or IgG as negative controls and β-actin as loading control (n = 2 experiments per receptor). Abbreviations: BP, blocking peptide; Ct, cycle threshold; IgG, immunoglobulin G; kDa, kilodalton; PC, positive control.

### ATP-Induced Ca²⁺ Signaling Reflects Only Intracellular Sources

Functionally active cell surface P2 receptors were identified by real-time Fura-2 calcium imaging in response to nucleotide stimulation. ATP (100 µM) rapidly induced a transient increase in cytosolic Ca²⁺ concentration ([Ca²⁺]_(cyt)_), which gradually returned to baseline (Fig. 2a). To determine the source of this Ca²⁺ signal, we performed ATP stimulation under Ca²⁺-free conditions and after endoplasmic reticulum (ER) store depletion with thapsigargin (10 µM). In Ca²⁺-free extracellular conditions, ATP still induced a Ca²⁺ signal, suggesting contribution from intracellular stores (Fig. 2b). Following ER depletion with thapsigargin, ATP failed to elicit a Ca²⁺ response, indicating that the ATP-induced Ca²⁺ rise primarily results from intracellular release (Fig. 2c). Δ_max_ (ATP-induced increase in [Ca²⁺]_(cyt)_, vs. baseline) was quantified per cell (Fig. 2d). hBECs showed Δ_max_ values of 101 ± 47 nM (Ca²⁺-containing), 84 ± 46 nM (Ca²⁺-free), and 0 ± 5 nM (thapsigargin, Ca²⁺-containing). A significant difference was observed only between ATP/Ca²⁺-containing and thapsigargin/ATP/Ca²⁺-containing conditions.

**Fig. 2.**
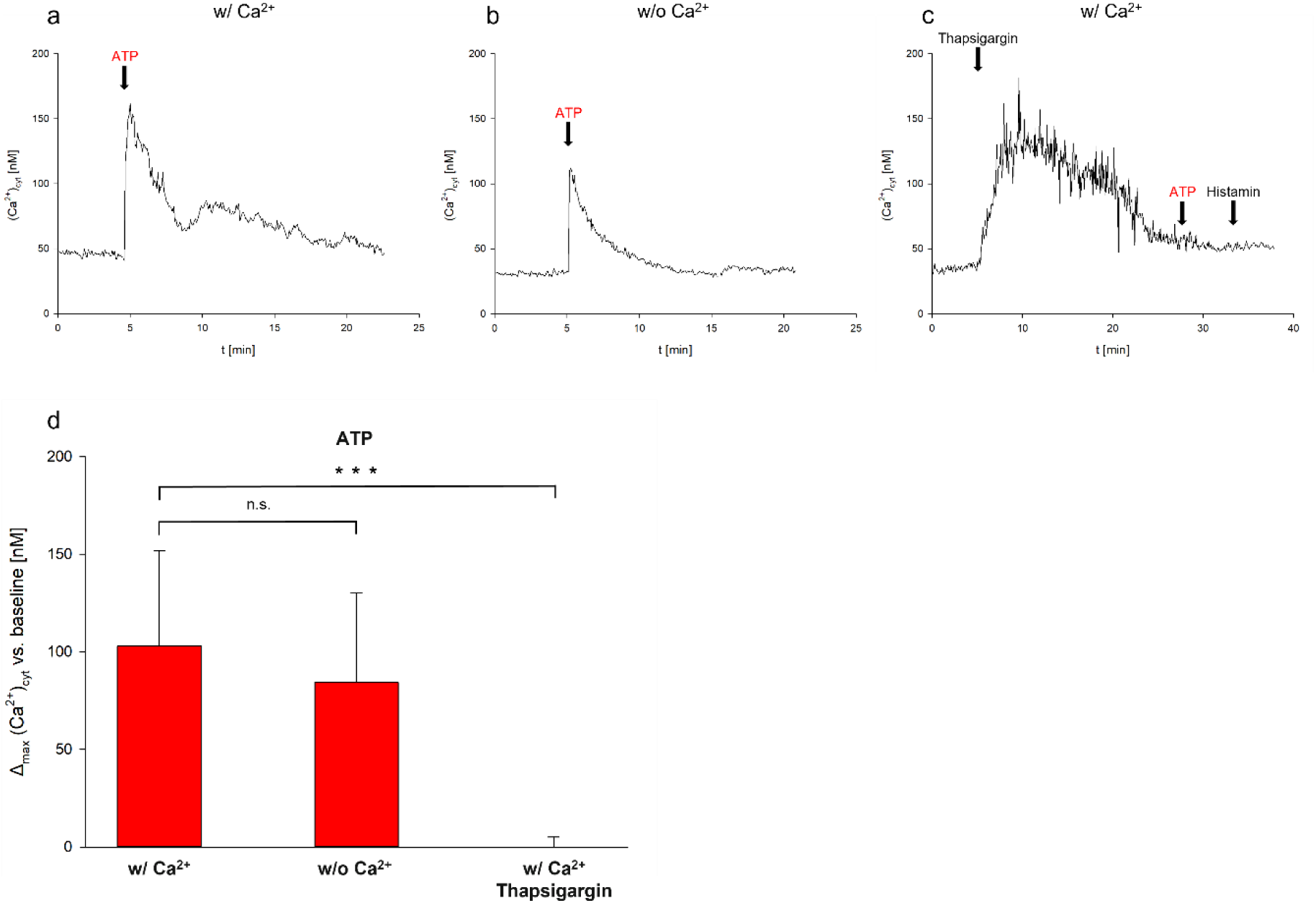
**a-c** [Ca^2+^]_(cyt)_ traces in representative hBECs after ATP (100 µM) under Ca^2+^-containing (**a**), Ca^2+^-free (**b**), or thapsigargin (10 µM, Ca^2+^ present) conditions (**c**); Histamine (100 µM) served as positive control. **d** Quantification of [Ca^2+^]_(cyt)_ responses (Δ_max_ vs. baseline) of a–c (w/ Ca^2+^, n = 13 experiments; w/o Ca^2+^, n = 11 experiments; w/ Ca^2+^ + thapsigargin, n = 10 experiments). *** p < 0.001, Mann–Whitney rank sum test. Abbreviations: cyt, cytosolic; nM, nanomolar; n.s., not significant; vs., versus; w/, with Ca^2+^; w/o, without Ca^2+^.

### Pharmacological Profiling Identifies P2Y_2_, P2Y_6_, and P2Y_12_ as Functionally Active Receptors

To investigate which of the expressed P2Y receptors were functionally coupled to intracellular signaling, we stimulated hBECs with selective natural or synthetic agonists (100 µM). UTP (P2Y_2_) induced robust Ca²⁺ signals, while UDP (P2Y_6_) and ADP (P2Y_12_) produced weaker, sustained responses (UTP > UDP > ADP) (Fig. 3a-c). In contrast, UDP-galactose (P2Y_14_ agonist) and α,β-methyleneATP (P2X_1/2/5/6_ agonist) failed to elicit Ca²⁺ signals, despite ATP responsiveness in the same cells (fig. 3d-e). Δ_max_ indicated concentrations of 5±3 nM for a,ß-meATP, 94±33 nM for UTP, 30±16 nM for UDP, 44±37 nM for ADP and 3±2 nM for UDP-galactose (fig. 3f). Statistical testing showed significant results only after stimulation with UTP, UDP, and ADP. These results indicate that P2Y_2_, P2Y_6_, and P2Y_12_ are functionally active in hBECs, with P2Y_2_ being the dominant receptor.

**Fig. 3.**
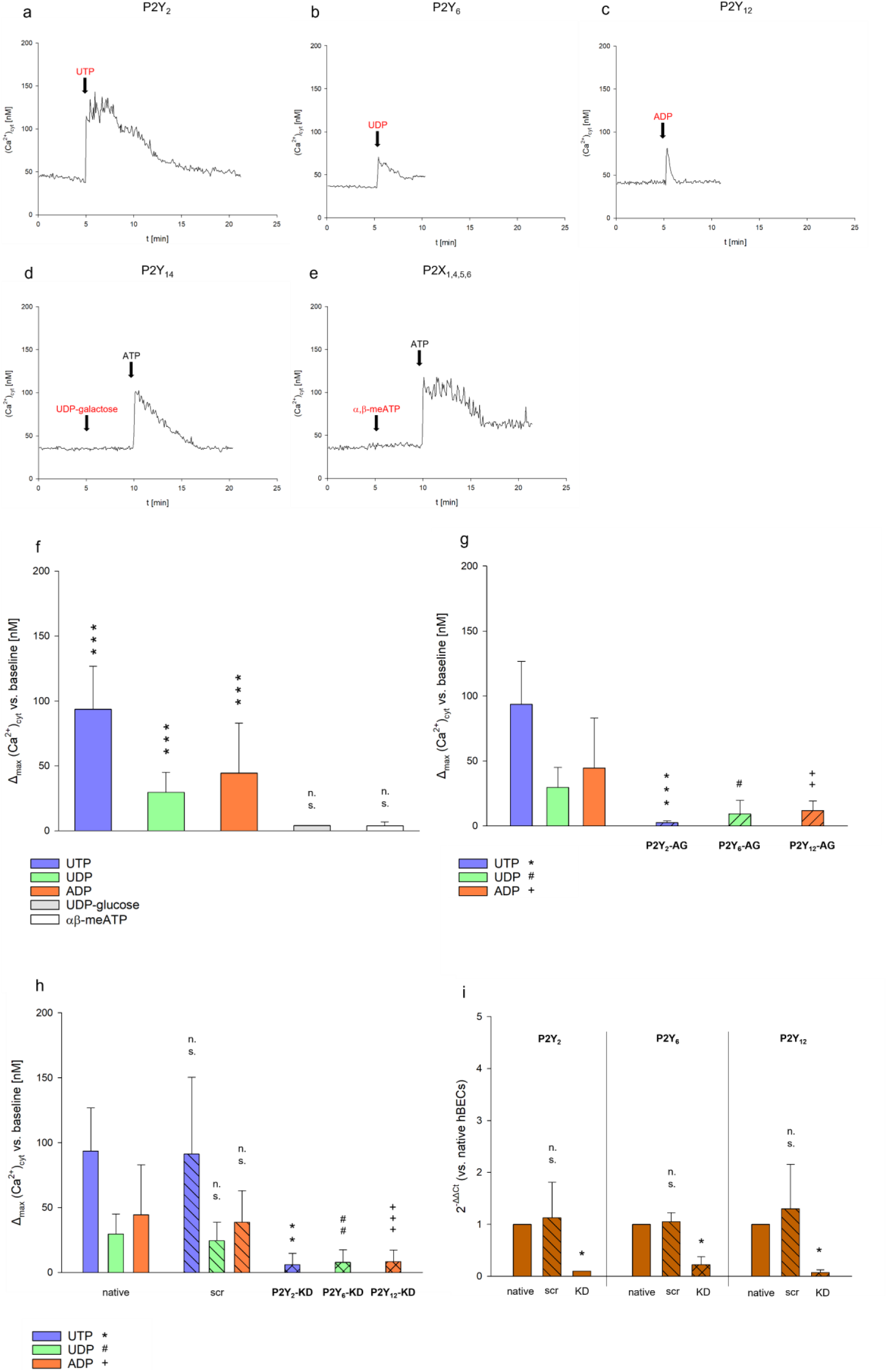
**a–e** [Ca^2+^]_(cyt)_ traces in representative hBECs after stimulation with UTP (**a**), UDP (**b**), ADP (**c**), UDP-galactose (**d**), or α,β-meATP (**e**) (each 100 µM); ATP (100 µM) served as positive control. **f** Quantification of [Ca^2+^]_(cyt)_ responses (Δ_max_ vs. baseline) of a-e: UTP (n = 9 experiments), UDP (n = 9 experiments), ADP (n = 11 v), UDP-galactose (n = 6 experiments), or α,β-meATP (n = 6 experiments). *** p < 0.001, Paired t-test (UTP, UDP) or Wilcoxon signed-rank test (ADP), intragroup comparison. **g-h** Quantification of [Ca^2+^]_(cyt)_ responses (Δ_max_ vs. baseline). **g** After UTP with P2Y_2_ antagonization (n = 7 experiments), UDP with P2Y_6_ antagonization (n = 6 experiments), or ADP with P2Y_12_ antagonization (n = 8 experiments), compared with native cells (f). # p < 0.05, ++ p < 0.01, *** p < 0.001, Student’s t-test (P2Y_2_, P2Y_6_ antagonists) or Mann–Whitney rank-sum test (P2Y_12_ antagonist), against native cells. **h** After UTP in P2Y_2_ knockdown (n = 11 experiments) or scramble (n = 13 experiments) cells, UDP in P2Y_6_ knockdown (n = 13 experiments) or scramble (n = 12 experiments) cells, or ADP in P2Y_12_ knockdown (n = 11 experiments) or scramble (n = 13 experiments) cells, compared with native cells (f). **/## p < 0.01, +++ p < 0.001, Mann–Whitney rank-sum test, scramble vs. knockdown. **i** mRNA expression of P2Y_2_, P2Y_6_, and P2Y_12_ in native, scramble, and knockdown hBECs assessed by RT-qPCR. Data are presented as 2^-^ΔΔCt relative to native hBECs, with RPL13A-1 as housekeeping gene (n = 4 experiments per receptor). * p < 0.05, Student’s t-test (P2Y_2_) or Mann–Whitney rank sum test (P2Y_12_, P2Y_6_), scramble vs. knockdown. Abbreviations: AG, antagonist; Ct, cycle threshold; cyt, cytosolic; KD, knockdown; nM, nanomolar; n.s., not significant; scr, scramble; vs., versus.

### Antagonism and Knockdown Confirm Receptor-Specific Ca^2+^ Responses

To verify receptor-specific Ca²⁺ responses, we performed pharmacological antagonism and siRNA-mediated knockdown of P2Y_2_, P2Y_6_, and P2Y_12_ receptors. Each antagonist—AR-C 118925XX (P2Y_2_), MRS 2578 (P2Y_6_), and PSB 0739 (P2Y_12_)—significantly blocked Ca²⁺ responses selectively to UTP (3±1 nM), UDP (9±11 nM), and ADP (12±7 nM), respectively (Fig. 3g). Similarly, receptor-specific siRNA knockdown significantly reduced the respective agonist-induced Ca²⁺ responses, yielding Δ_max_ values of 6 ± 9 nM for UTP, 8 ± 10 nM for UDP, and 8 ± 9 nM for ADP, whereas control siRNA-treated cells maintained normal signaling (UTP 91±59, UDP 24±14, ADP 39±24) (Fig. 3h). RT-qPCR confirmed efficient and specific knockdown for each target receptor (Fig. 3i).

### P2Y_2_ Signaling Licenses Functional Activation of P2Y_6_ and P2Y_12_

To explore possible cross-regulation between purinergic receptors, we examined whether prior activation, inhibition, or absence of P2Y_2_ affects signaling via P2Y_6_ and P2Y_12_. Strikingly, pre-stimulation with UTP significantly abolished subsequent UDP- (3±3 nM) and ADP-induced Ca²⁺ responses (7±3 nM) as well as pharmacological inhibition of P2Y_2_ (UDP 3±1 nM, ADP 3±2 nM) (Fig. 4a). Similarly, genetic knockdown of P2Y_2_ also significantly suppressed responses to both UDP (3±2 nM) and ADP (5±6 nM) (Fig. 4b). These effects were not due to downregulation of P2Y_6_ or P2Y_12_ expression, as transcript levels remained stable or were slightly elevated following P2Y_2_ knockdown (Fig. 4c). These data suggest that P2Y_2_ expression is necessary for downstream P2Y_6_ and P2Y_12_ signaling while upstream activation of P2Y_2_ impairs function of P2Y_6_ and P2Y_12_ — consistent with a hierarchical, gatekeeping role for P2Y_2_ in purinergic signaling.

**Fig. 4.**
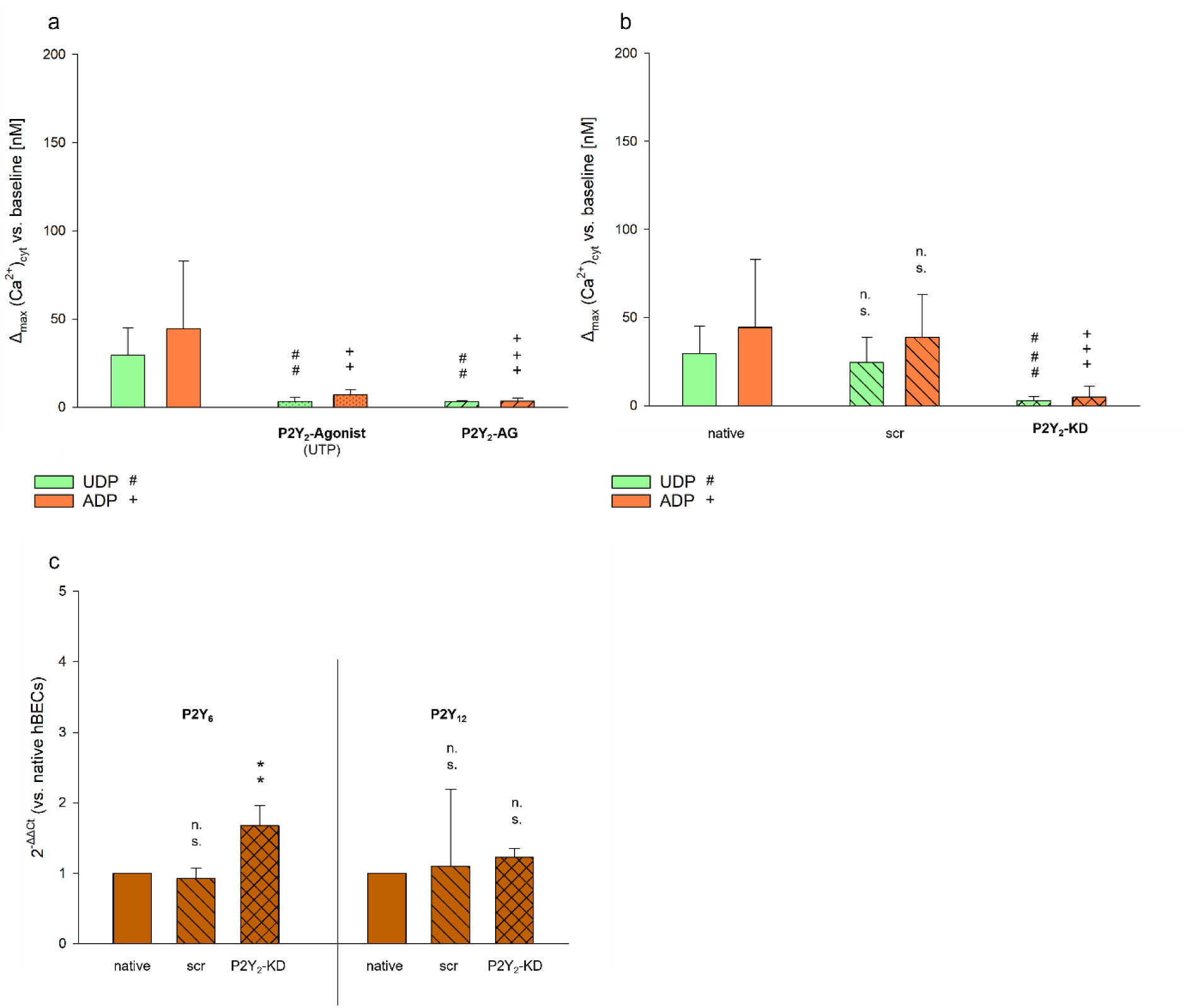
**a-b** Quantification of [Ca^2+^]_(cyt)_ responses (Δ_max_ vs. baseline) in hBECs. **a** After stimulation with UDP or ADP (each 100 µM) following primary stimulation with UTP (100 µM) (n = 6 experiments per agonist), or after P2Y_2_ antagonization (n = 6 experiments per agonist), compared with native cells (3f). ##/++ p < 0.01, +++ p < 0.001, Student’s t-test (UDP) or Mann–Whitney rank-sum test (ADP), against native cells. **b** After stimulation with UDP or ADP (each 100 µM) in P2Y_2_ knockdown cells (UDP, n = 10; ADP, n= 11 experiments), compared with scramble and native cells (3h). ###/+++ p < 0.001, Student’s t-test (UDP) or Mann–Whitney rank-sum test (ADP), scramble vs. knockdown. **c** Expression of P2Y_6_ and P2Y_12_ at the mRNA level in hBECs after P2Y_2_ knockdown, compared with native or scramble cells, assessed by RT-qPCR. Expression shown as 2^-^ΔΔCt relative to native hBECs, with RPL13A-1 as housekeeping gene (n = 4 experiments per receptor). ** p < 0.01, Student’s t-test, scramble vs. knockdown. Abbreviations: AG, antagonist; Ct, cycle threshold; cyt, cytosolic; KD, knockdown; nM, nanomolar; n.s., not significant; scr, scramble; vs., versus

### Validation of Receptor Function in ALI-Cultured hBECs

To confirm the physiological relevance of our findings, we repeated key experiments in hBECs cultured at the air-liquid interface (ALI) for ∼28 days to mimic in vivo airway differentiation (Fig. 5a). ATP stimulation evoked Ca²⁺ responses (w/ Ca^2+^ 40±10 nM , w/o Ca^2+^ 53±9 nM) comparable to submerged cultures and was abolished after thapsigargin treatment (8±3 nM), confirming dependence on intracellular stores (Fig. 5b). UTP (54±30 nM), UDP (39±23 nM), and ADP (37±15 nM) all induced Ca²⁺ responses, comparable to submerged conditions, while α,β-methyleneATP again failed to do so (7±8 nM). Unexpectedly, UDP-galactose induced a significant calcium response (15±13 nM), solely mediated by the P2Y₁₄ receptor (5c). Antagonizing of the P2Y_2_ and P2Y_12_ again significantly abolished the UTP- (2±5 nM) and ADP-induced responses (8±6 nM), while in contrast to submerged conditions, UDP signaling was preserved after antagonizing the P2Y_6_ receptor (25±10nM), (Fig. 5d). Due to the lack of expression control and knockdown tools under ALI conditions, we applied highly selective agonists for P2Y₁, P2Y₄, P2Y₁₁, and P2Y₁₃ to unambiguously attribute the UTP-, UDP-, and ADP-induced signals to P2Y₂, P2Y₆, and P2Y₁₂, thereby ruling out misinterpretation. None of the control agonists elicited consistent calcium responses (P2Y₁ 4±2 nM, P2Y₄ 6±3 nM, P2Y₁₁ 9±9 nM, and P2Y₁₃ 3±2 nM); however, the P2Y₁ agonist reached statistical significance (Fig. 5e). As in submerged cultures, prior P2Y_2_ stimulation significantly eliminated Ca²⁺ responses to UDP (4±3 nM) and ADP (6±8 nM) as well as pharmacological inhibition of the P2Y_2_ receptor (UDP 3±6 nM, ADP 4±6 nM) (Fig. 6a).

**Fig. 5.**
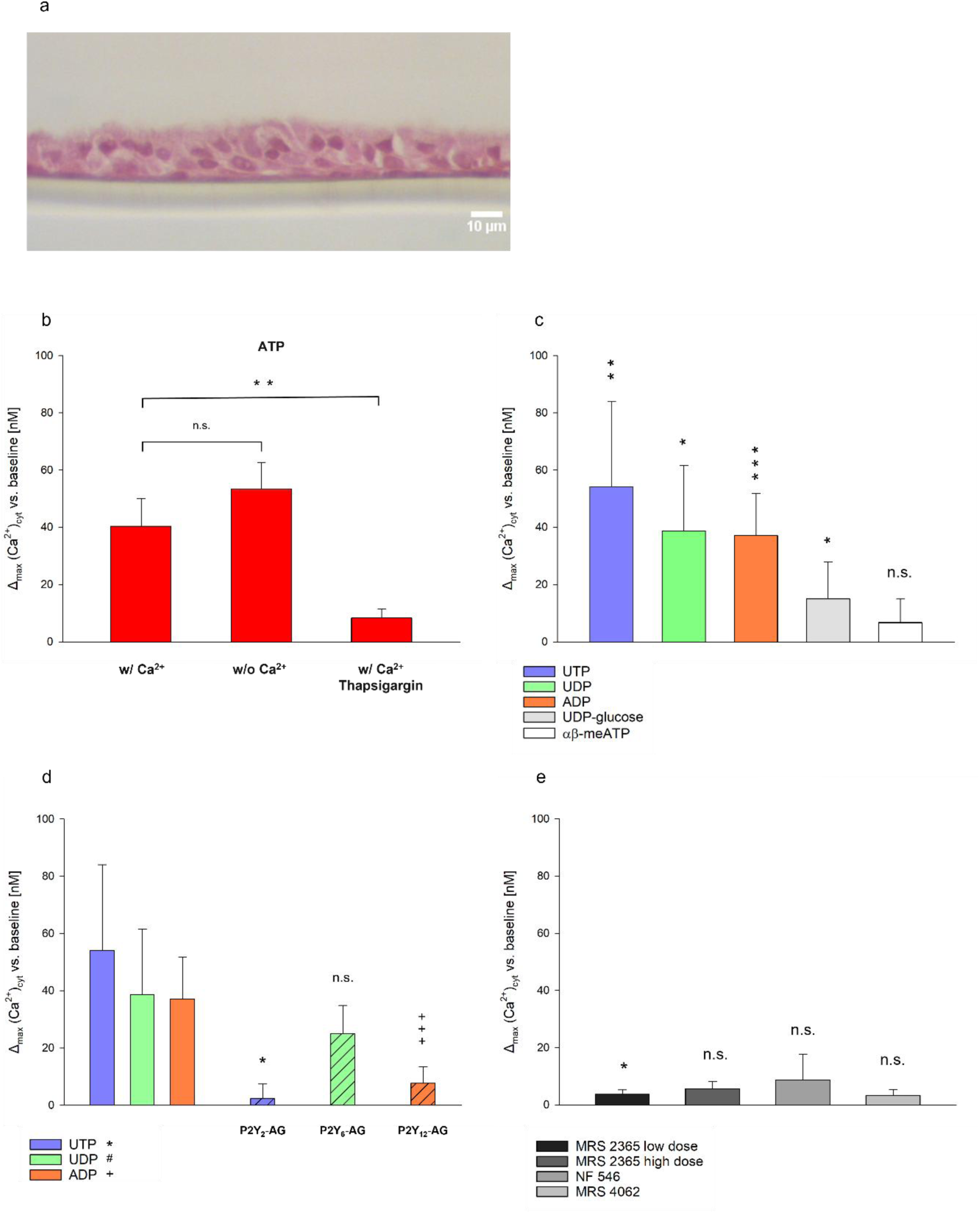
**a** Representative image of a paraffin-embedded section of a 3D-ALI model on a porous membrane after H&E staining. A multilayered epithelium with apical cilia is visible. **b–e** Quantification of [Ca^2+^]_(cyt)_ responses (Δ_max_ vs. baseline) in 3D-hBECs. **b** After apical ATP stimulation (100 µM) under Ca^2+^-containing, Ca^2+^-free, or thapsigargin (10 µM, Ca^2+^ present) conditions (n = 3 experiments per group). **p < 0.001, Student’s t-test. **c** After apical stimulation with UTP (n =6 experiments), UDP (n =6 experiments), ADP (n =7 experiments), UDP-galactose (n =7 experiments), or α,β-meATP (n =7 experiments) (each 100 µM). * p < 0.05, ** p < 0.01, *** p < 0.001, paired t-test (UTP, ADP) or Wilcoxon signed-rank test (UDP, UDP-galactose), intragroup comparison. **d** After apical stimulation with MRS2365 (5 nM, P2Y_1_ agonist,), MRS2365 (1 µM, P2Y_13_ agonist), NF546 (1 nM, P2Y_11_ agonist), or MRS4062 (50 nM, P2Y_4_ agonist) (n = 3 experiments per agonist). * p < 0.05, paired t-test, intragroup comparison. **e** After apical stimulation with UTP with P2Y_2_ antagonization (n = 3 experiments), UDP with P2Y_6_ antagonization (n = 7 experiments) or ADP with P2Y_12_ antagonization (n = 6 experiments), compared with native cells (c). * p < 0.05, +++ p < 0.001; Student’s t-test, against native cells. Abbreviations: AG, antagonist; cyt, cytosolic; nM, nanomolar; n.s., not significant; vs., versus.

**Fig. 6.**
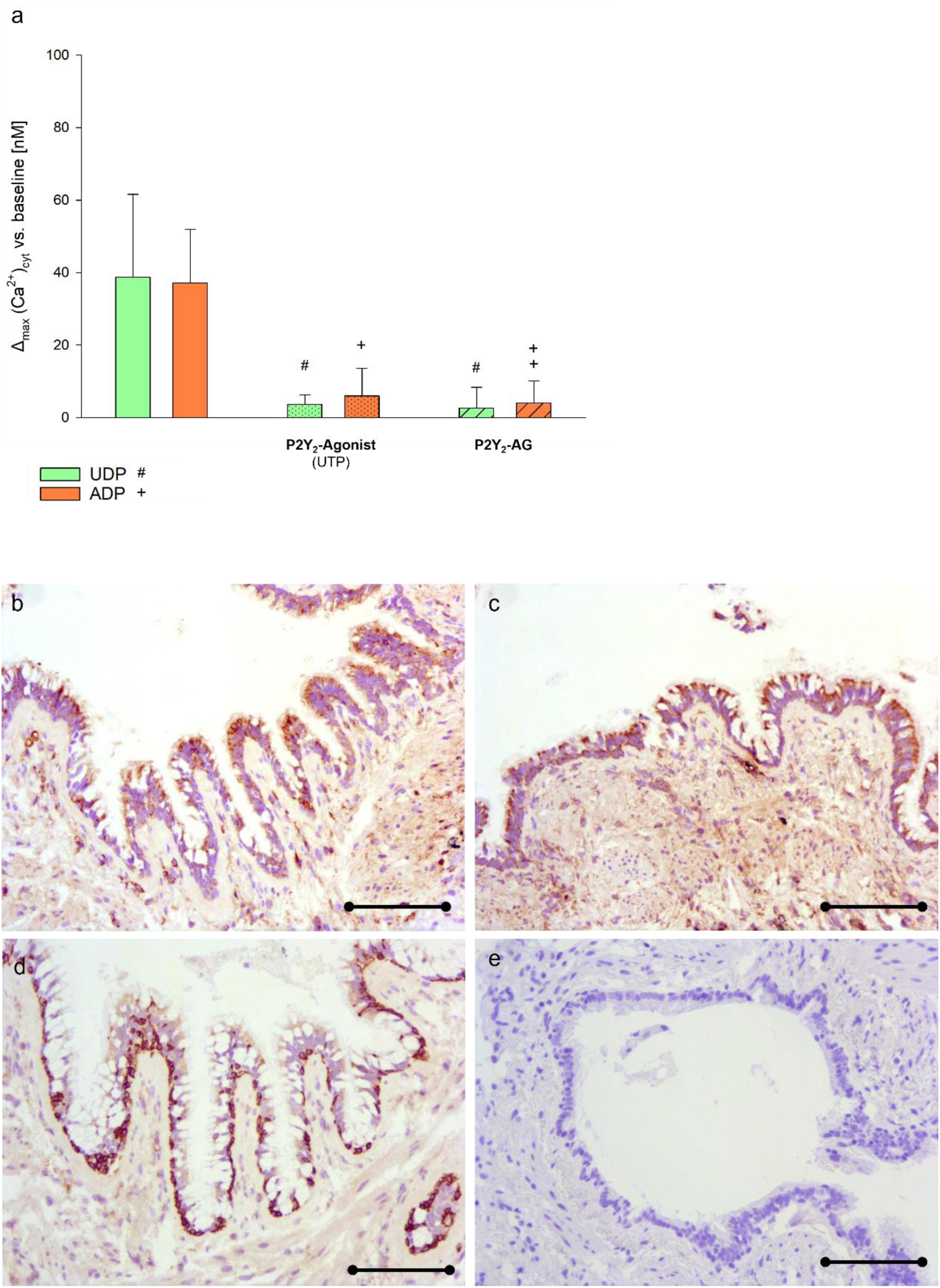
**a** Quantification of [Ca^2+^]_(cyt)_ responses (Δ_max_ vs. baseline) in 3D-hBECs after stimulation with UDP or ADP (100 µM) following primary stimulation with UTP (100 µM) (n = 3 experiments per agonist) or after P2Y_2_ antagonization (n = 3 experiments per agonist) compared with native cells (n=5c). #/+ p<0.05, ++ p< 0.01, Student’s t-test, against native cells. **b–e** Representative immunostaining of human bronchial sections with primary antibodies against P2Y_2_ (**b**), the P2Y_6_ (**c**) and the P2Y_12_ (**d**) receptor. Cells were counterstained with H&E. Brown coloration indicates receptor localization. IgG negative control with H&E staining (**e**). Abbreviations: AG, antagonist; cyt, cytosolic; nM, nanomolar; vs., versus.

### P2Y_2_, P2Y_6_, and P2Y_12_ Are Co-Expressed in Human Airway Tissue

To evaluate the in vivo relevance of our findings, we performed immunohistochemical staining for P2Y_2_, P2Y_6_, and P2Y_12_ in human bronchiolar biopsy specimens. P2Y_2_ and P2Y_6_ were found on both apical and basolateral sides of the airway epithelium, with higher intensity on the apical surface (Fig. 6 b–d). In contrast, P2Y_12_ was almost exclusively localized to the basolateral domain (Fig. 6 e). No signal was detected in IgG controls (Fig. 6E). These results support the physiological co-expression and spatial segregation of these receptors in the human airways.

## Discussion

Our study reveals a previously unrecognized hierarchical organization of purinergic signaling in hBECs, wherein the P2Y_2_ receptor serves not only as a primary responder to extracellular ATP or UTP but also as a necessary gatekeeper for the downstream activation of P2Y_6_ and P2Y_12_ receptors. This conditional relationship challenges again the canonical view of P2 receptors as parallel and independent conduits and suggests a logic-based architecture of signal integration in airway epithelial cells. Our findings highlight both the mechanistic sophistication of purinergic signaling and the importance of receptor cross-talk in maintaining specificity in a system exposed to rapidly fluctuating nucleotide concentrations.

### P2Y_2_ as a Central Integrator of Epithelial Purinergic Signaling

ATP, as a damage-associated molecular pattern, is rapidly released from epithelial cells during mechanical stress, infection, or inflammation [23]. It serves as a key signal for initiating repair and immune responses [24]. P2Y₂ is consistently acknowledged as the key P2 receptor in airway physiology, especially in mediating ATP-driven responses, yet the basis of its functional dominance has remained largely unexplored [25, 26]. In our model, P2Y₂ serves as a molecular integrator, linking ATP/UTP-induced calcium signaling to the suppression of P2Y₆ and P2Y₁₂ activity, thereby elevating its functional significance.

### Receptor Interdependence and Potential for Heteromerization

Mechanistically, we show that siRNA-mediated knockdown of P2Y_2_ abolishes calcium responses to both UDP and ADP, despite intrinsic expression of their respective receptors, P2Y_6_ and P2Y_12_. These findings attribute a permissive role to P2Y_2_, as its expression is an essential prerequisite for P2Y₆ and P2Y₁₂ function in our cell model. Notably, transcript levels of P2Y_6_ and P2Y_12_ were unaffected or slightly increased following P2Y_2_ knockdown, indicating that the permissive effect of P2Y₂ is mediated through post-transcriptional mechanisms—such as post-translational modifications, membrane compartmentalization, or direct receptor–receptor interactions [27].

The functional dependency between the NTP-sensitive receptor P2Y_2_ and the co-expressed NDP-sensitive receptors, P2Y_6_ and P2Y_12_, was one of the most striking findings. Prior pharmacological activation or blockade of P2Y₂ consistently suppressed downstream responses mediated by P2Y₆ and P2Y₁₂, implicating a coordinated mechanism that prevents accidental signaling through secondary nucleotides. This was observed in both submerged and 3D ALI cultures, indicating the robustness of this regulatory mechanism across epithelial states. The observed functional dependency may be structurally mediated, given that the P2Y₂ antagonist did not trigger a calcium response on its own, arguing against mechanisms involving intracellular Ca²⁺ depletion, second messenger competition, or heterologous desensitization [28]. Although our preliminary co-immunoprecipitation experiments were inconclusive, the functional loss of P2Y_6_ and P2Y_12_ signaling following P2Y_2_ knockdown — without changes in receptor expression — further suggests potential receptor complex formation, such as heteromers, or shared scaffolding components required for proper function.

Interestingly, P2Y_12_ — a G_i_ coupled receptor not previously reported in the lower human airway epithelium [29, 30]— was confirmed at the mRNA and protein levels using enhanced RT-qPCR, western blot and immunohistochemistry. Selective agonists and siRNA validated its functional role in ADP-mediated Ca²⁺ signaling, suggesting a novel site of action that may influence immune cell recruitment, platelet-like signaling, or epithelial-stromal communication from the basolateral epithelial surface, where it is specifically localized. Notably, during the preparation of this manuscript, evidence for P2Y_12_ expression in an immortalized human bronchial epithelial cell line was reported, but not validated at the protein level or in a physiological model [31].

### Physiological and Spatial Context: Licensing as a Protective Filter

This hierarchical gating model serves an important physiological function: it may act as a biological filter, ensuring that UDP and ADP — which are more abundant and less specific signals due to their derivation from ATP — do not trigger immune or secretory responses unless they are primarily released. Given that ecto-nucleotidases such as NTPDase rapidly degrade ATP and UTP to ADP and UDP in the extracellular space [1, 2], the system is inherently prone to background noise. P2Y_2_ may have evolved as a spatial and temporal checkpoint that imposes order on a potentially noisy signal landscape in adaption to environmental changes.

The importance of this biological filter is highlighted by the permissive role of P2Y₂, which prevents the functional co-existence of NTP- and NDP-sensitive receptors outside this hierarchical cluster in our cellular model.

This is supported by spatial expression patterns observed in human bronchial biopsies: P2Y_2_ and P2Y_6_ were detected apically and basolaterally, while P2Y_12_ was confined to the basolateral membrane, likely linking it to stromal or immune interactions. The combination of receptor co-expression and spatial compartmentalization strengthens the model of localized, regulated activation of downstream pathways, particularly in the context of mucosal immunity.

### Model Systems and Functional Insights

Our findings also demonstrate the importance of choosing appropriate model systems. While submerged hBEC cultures provided clarity for mechanistic dissection, 3D ALI cultures more closely recapitulated in vivo conditions, enabling detection of P2Y_14_ receptor activity, as suggested by UDP-galactose–induced Ca²⁺ signals. P2X receptor function remained limited in both systems, despite detectable mRNA and protein expression, possibly reflecting their known conditional activation during inflammation or infection. Additionally, 3D cultures revealed limitations in antagonist accessibility and reproducibility — for example, the failure of P2Y_6_ antagonists to block UDP responses under ALI conditions — highlighting the influence of the epithelial matrix, mucus, or cilia on receptor pharmacology [32].

### Limitations and Future Directions

Several questions remain. What is the molecular basis of the observed receptor hierarchy? Does P2Y_2_ form heteromeric complexes with P2Y_6_ or P2Y_12_, or does it prime the signaling environment via Ca²⁺-dependent second messengers or membrane microdomains? Can this licensing model be generalized to other epithelial tissues or disease states? Future studies should include super-resolution imaging, co-IP validation, proximity ligation assays, and analyses of downstream signaling branches beyond calcium, such as MAPK, NF-κB, or cytokine secretion. The functional consequences of P2Y_12_ activity in epithelial immunity and its relevance in smoking-related or inflammatory airway diseases also warrant deeper investigation [33].

## Conclusion

Taken together, our findings support a new model of hierarchical, conditional purinergic signaling in the human airway epithelium, where P2Y_2_ serves as a central integrator that governs access to downstream pathways mediated by P2Y_6_ and P2Y_12_. This architecture imposes a temporal logic on epithelial responses to extracellular nucleotides, enabling refined control of mucosal defense, repair, and homeostasis. It also provides a framework for therapeutic strategies targeting receptor interdependencies rather than individual receptors in isolation — an approach that may prove valuable in diseases characterized by purinergic dysregulation, such as asthma, COPD, or cystic fibrosis.

## Methods

### Isolation of Primary Human Bronchial Epithelial Cells

Primary human bronchial epithelial cells were isolated from residual donor lung tissue following partial lung resection at the Department of General, Visceral and Thoracic Surgery, University Medical Center Hamburg-Eppendorf, Germany. Ethical approval was obtained from the Medical Council of Hamburg. Pulmonary lobes were immediately immersed in Ringer’s lactate solution (B. Braun Melsungen AG, Germany) at 4°C and processed within 4 hours.

Bronchi from segmental and subsegmental airways were dissected under semi-sterile conditions, cut into 1–2 cm segments, and then longitudinally bisected. Tissue pieces were placed in Petri dishes (Greiner Bio-One) with cold PBS and kept at 4°C. All subsequent steps were performed under a sterile laminar flow hood.

Tissue was washed twice in PBS and incubated in 25 mL of Wash Solution A (tab. 1) for 30 min at 4°C. After two washes in DMEM, segments were transferred into Wash Solution B for 48 h at 4°C on a shaker, with the solution renewed after 24 h. Following this, tissue was briefly equilibrated in RPMI supplemented with 10% FBS for 5 min at room temperature.

**Tab. 1.** Wash solutions A and B, as well as Starter Medium, were used for the isolation and early-stage cultivation of the primary cells.

| Wash solution A | Wash solution B | Starter medium |
| --- | --- | --- |
| DMEM + DNase 10 µg/ml + dithiothreitol 500 µg/ml + gentamicin 50 µg/ml + amphotericin 0,25 µg/ml + 1% pen/strep | DMEM + DNase 10 g/ml + gentamicin 50 µg/ml + amphotericin 0,25 µg/ml + protease 0,05% + 1% pen/strep | AEGM + gentamicin 50 µg/ml + amphotericin 0,25 µg/ml + 1% pen/strep |

Epithelial cells were collected by scraping the luminal surface with a sterile scalpel and rinsing with RPMI + 10% FBS. The cell suspension was centrifuged at 200 × g for 5 min, and the pellet was resuspended in 3 mL starter medium (Table 1). Cells were cultured in T25 flasks (Greiner) with 6 mL medium at 37°C, 5% CO₂. The medium was replaced every other day. After 7 days, the culture was switched to Airway Epithelial Growth Medium (AEGM) supplemented with 10,000 U/mL penicillin-streptomycin.

Upon reaching 90% confluence, cells were washed with PBS, detached using Accutase (2 mL, 6 min, 37°C), and centrifuged. The pellet was resuspended in AEGM and transferred to T75 flasks for expansion.

### Cell Culture Conditions

#### Submerged Culture

Cells were maintained in T75 flasks in 12 mL AEGM with 10,000 U/mL penicillin-streptomycin at 37°C, 5% CO₂. For subculturing, cells at 90% confluence were washed twice with PBS, incubated with Accutase, and reseeded at a 1:3 ratio. Mycoplasma contamination was routinely ruled out via PCR.

#### Air-Liquid Interface (ALI) Culture

Commercial primary bronchial epithelial cells (PromoCell, Heidelberg, Germany), pre-screened for their ability to form tight epithelial layers, were used for ALI culture. After expansion under submerged conditions, cells were seeded onto 24-well Transwell inserts (0.4 μm pore size, CellQuart), pre-coated with collagen (Merck), in AEGM with 1% penicillin-streptomycin applied to both apical and basolateral chambers.

After ∼3 days, once full confluence and a rise in transepithelial electrical resistance (TEER) were confirmed by microscopy and TEER measurement, medium was removed from the apical side and replaced basolaterally with ALI-Airway Medium (PromoCell) supplemented with 50 μg/mL gentamicin. Medium was refreshed every 2–3 days, and cells were washed weekly with PBS on both sides. TEER was measured weekly using a Millicell-ERS2 Volt-Ohm Meter (MilliporeSigma) with STX01 electrodes. Cultures with insufficient or declining TEER were excluded.

#### Immunofluorescence Characterization

To confirm epithelial identity, cytokeratin 18 expression was assessed in cells subcultured three times. Cells (5 × 10⁴) were seeded on Falcon™ 4-well chamber slides (Thermo Fisher) and cultured for 48 h. After one PBS wash, chambers were removed, and cells were fixed in cold methanol:acetone (7:3) for 10 min at −20°C.

Following three PBS washes, samples were blocked with 1% BSA (Thermo Fisher) for 1 h at room temperature and incubated with either anti-cytokeratin 18 or isotype control antibodies (Abcam) for 1 h at 37°C (tab. 2). After three washes, cells were incubated with Alexa Fluor 488-conjugated secondary antibodies for 1 h. Nuclei were stained with Hoechst 33342 (0.5 µg/mL) for 10 min, and slides were mounted with Fluoromount-G. Imaging was performed using an epifluorescence microscope (Olympus) with filters for Alexa Fluor 488 and Hoechst (Fig. S1.).

**Tab. 2.** Antibodies and isotype controls applied for the detection of cytokeratin 18 in hBECs by immunocytochemistry.

| Primary antibody | Isotype control | Secondary antibody |
| --- | --- | --- |
| Anti-cytokeratin 18, 1:100 diluted in PBS | mIgG1 [B11/6] - isotype control, 1:100 diluted in PBS | Goat anti-mIgG H&L Alexa Fluor® 488, 1:200 diluted in PBS |

### Intracellular Ca²⁺ Imaging

#### Submerged Culture

Cells were seeded on collagen-coated glass-bottom dishes (Corning) and incubated for 48 h. Cells were loaded with 10 µM fura-2-AM in HBSS (pH 7.3) for 40 min at 37°C in the dark. Intracellular Ca²⁺ was measured by dual-excitation ratiometric imaging (340/380 nm) on an epifluorescence microscope (Olympus) with MetaFluor software (Molecular Devices).

Ca²⁺ signals were stimulated with 100 µM nucleotides (ATP, UTP, ADP, UDP, α,β-methyleneATP, UDP-galactose). ATP and histamine (100 µM) served as positive controls. P2 receptor antagonists (10 µM) were added 15 min prior to stimulation where indicated.

#### ALI Culture

For ALI cultures, fura-2-AM loading was performed from both apical and basolateral sides. Inserts were carefully removed and mounted on collagen-coated glass-bottom dishes using beeswax at the edges, avoiding contact with the cell layer. Cells were covered with HBSS, and Ca²⁺ imaging was performed as described.

### Quantitative RT-PCR

Total RNA was extracted using the RNeasy® Plus Mini Kit (Qiagen), and concentrations were measured using a NanoDrop spectrophotometer. cDNA synthesis was performed using the Maxima First Strand Kit (Thermo Fisher). For low-abundance transcripts such as P2Y12, preamplification was performed (PrimePCR PreAmp, Bio-Rad). qPCR was performed using gene-specific primers (tab. 3) designed to span exon-exon junctions. Reactions used QuantiNova SYBR Green (Qiagen), except for P2Y12, which used PrimePCR SYBR Green (Bio-Rad). Runs included negative and no-template controls. RPL13A served as a reference gene. Data were analyzed using the ΔΔCt method, with native hBECs as the reference.

**Tab. 3.**
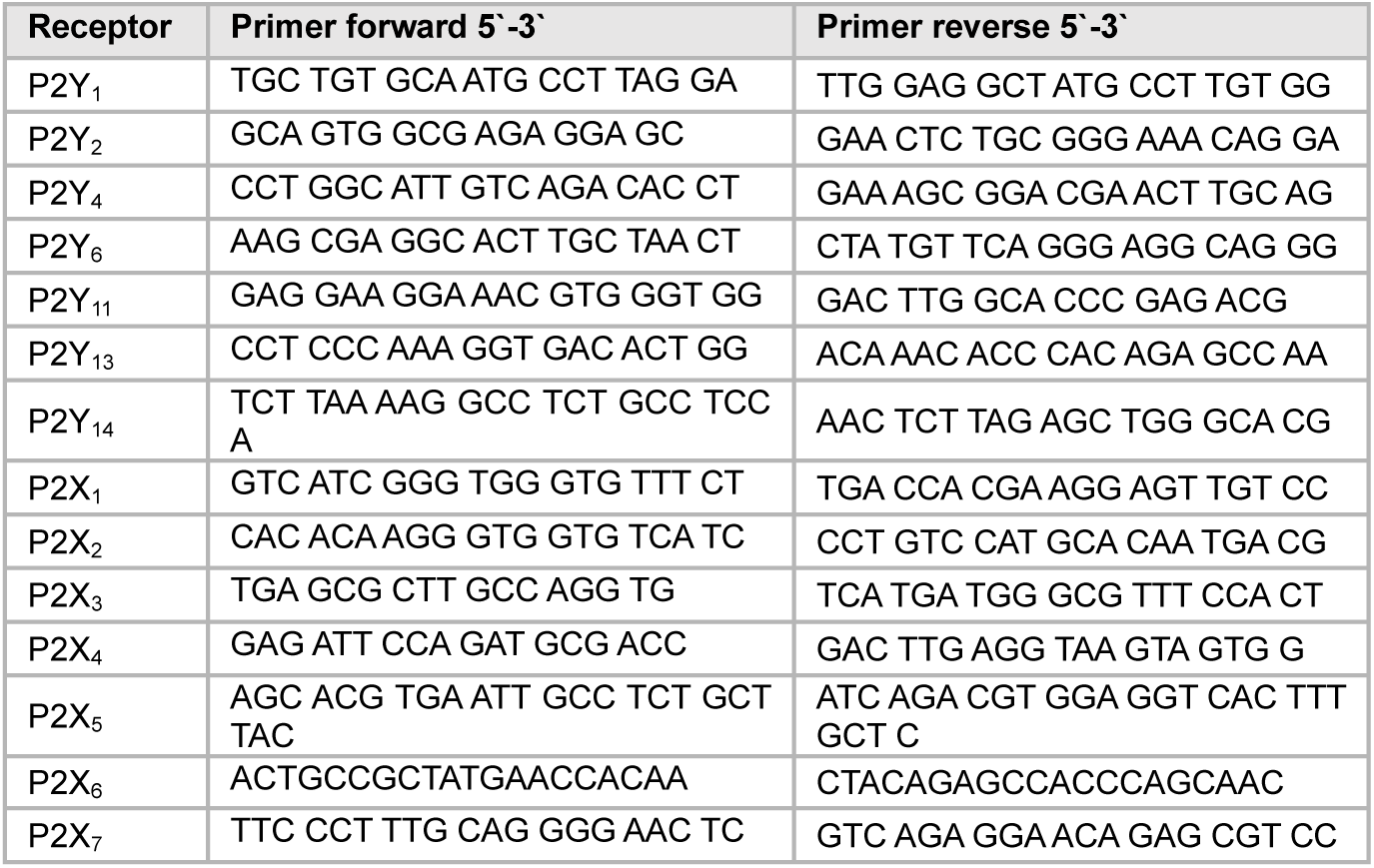
The primer pairs for P2R detection were designed using the National Center for Biotechnology Information (NCBI) database and synthesized by Eurofins.

### Western Blotting

Cells were lysed in RIPA buffer with protease inhibitors (Roche), and protein content was quantified by BCA assay. Samples were denatured at 95°C and separated by SDS-PAGE (4–12% Bis-Tris gels, Life Technologies), then transferred to nitrocellulose membranes.

Membranes were blocked in 5% milk in PBS-T and incubated with primary antibodies against P2X and P2Y receptors (tab. 4) overnight at 4°C. HRP-conjugated secondary antibodies were applied for 1 h at room temperature, and bands were visualized using ECL (GE Healthcare). β-actin was used as a loading control. Antibody specificity was confirmed with blocking peptides or isotype controls.

**Tab. 4.** Primary antibodies with epitopes and dilutions used in WB experiments.

| Receptor | Epitop | Dilution | Brand |
| --- | --- | --- | --- |
| P2Y<sub>1</sub> | (unknown) 19 amino acid peptide from C-terminus | 1:200 | Genway |
| P2Y<sub>2</sub> | 227-244 | 1:400 | Alomone |
| P2Y<sub>4</sub> | 170-220 | 1:200 | Santa Cruz |
| P2Y<sub>6</sub> | 311-328 | 1:200 | Alomone |
| P2Y<sub>11</sub> | (unknown) 17 amino acid peptide from 3rd cytoplasmic domain | 1:400 | Genway |
| P2Y<sub>12</sub> | 125-142 | 1:200 | Alomone |
| P2Y<sub>13</sub> | 119-134 | 1:200 | Alomone |
| P2Y<sub>14</sub> | 256-305 | 1:1000 | Abcam |
| P2X<sub>1</sub> | 382-399 | 1:200 | Alomone |
| P2X<sub>2</sub> | 457-472 | 1:200 | Alomone |
| P2X<sub>3</sub> | 383-397 | 1:200 | Alomone |
| P2X<sub>4</sub> | 370-388 | 1:200 | Alomone |
| P2X<sub>5</sub> | 437-455 | 1:200 | Alomone |
| P2X<sub>6</sub> | 363-379 | 1:200 | Alomone |
| P2X<sub>7</sub> | 576-595 | 1:500 | Alomone |

### siRNA-Mediated Knockdown

Cells were seeded in antibiotic-free medium and transfected with 10 nM siRNA targeting P2Y2, P2Y6, or P2Y12 (Qiagen) using Lipofectamine RNAiMAX (Thermo Fisher). After 6 h, transfection medium was replaced. After 48 h, cells were used for RT-qPCR or Ca²⁺ imaging. Knockdown efficiency was confirmed by qPCR.

### Immunohistochemistry

Human lung tissues were obtained from surgical resections due to lung cancer, provided by the Biomaterialbank North, and ethical consent was obtained from the ethics commission of the University of Lübeck (AZ 12-220). Tissues at least 10 cm distant from the tumors were used in this study.

Hope-fixed, paraffin-embedded lung tissues were prepared as previously described [34]. Sections of 4 µm were cut, and the slides were deparaffinized. Primary antibodies (the same as used for western blot experiments in hBECs) were applied at a dilution of 1:400 in antibody diluent (Zytomed Systems, Berlin). Detection was achieved using the ZytochemPlus HRP polymer kit (Zytomed Systems, Berlin) with 30 min incubation, and color development was performed using permanent aminoethylcarbazole (AEC, Zytomed Systems, Berlin) as the chromogen (10 min). Slides were counterstained with Mayer’s hemalum and mounted. Negative controls were included in every staining series. Slides were photographed using a Nikon Eclipse 80i at 400× magnification.

### Histology of ALI Cultures

Membranes with 28-day ALI cultures were fixed in 3.5% formalin (BÜFA Chemikalien) for 24 h at 2°C, embedded in paraffin, and sectioned at 5 µm. Sections were mounted, deparaffinized, stained with hematoxylin and eosin, and coverslipped with Eukitt (Merck). Morphology was assessed by light microscopy (Leica).

### Data Analysis

Ca²⁺ imaging results are presented as mean ± SD. Statistical analyses were performed using SigmaStat (Systat Software) employing Two-tailed Student’s t-tests, Mann–Whitney rank sum tests, Paired t-test or Wilcoxon signed-rank tests as appropriate. Significance was set at *p* < 0.05.

RT-qPCR data were normalized to RPL13A, and expression values are shown as mean ± SD. Expression profiling of P2 receptors (yes/no screening) was not statistically analyzed. Knockdown efficiency was evaluated using the ΔΔCt method.

### Language editing

During the preparation of this work the author used ChatGPT (OpenAI, San Francisco, CA, USA) in order to improve language editing. The tool was used exclusively to improve grammar and clarity; it was not involved in generating or altering the scientific content. All final edits were reviewed and approved by the authors.

## Acknowledgment

The authors wish to thank Claudia Luechau, Christiane Pahrmann, Andrea Pawelczyk, Kirsten Pfeiffer-Drenkhahn, Sabine Vidal-y-Syand, Monika Weber and Ewa Wladykowski for their technical assistance and dedicated work on this project. Furthermore, they wish to thank Sonja Schrepfer for providing the lung tissue for isolation of bronchial epithelial cells.

## Author contributions

Volker Meidl: study design, performance of experiments, analysis and interpretation of data, drafting of the manuscript.

Christian Börnchen: Performance of experiments.

Martina Kiefmann and Rainer Kiefmann: study design, interpretation of the data and revision of the manuscript.

Torsten Goldmann: Performance and interpretation of immunohistochemistry experiments of human bronchial biopsies.

## Materials & Correspondence

Volker Meidl

