## Supplemental Figure S1 for "A Hierarchical Model of Purinergic Receptor Activation in Bronchial Epithelial Cells"

**Supplementary material**

Fig. S1: Immunocytochemistry demonstrates clear expression of cytokeratin 18 in primary isolated cells. The absence of fluorescence following incubation with the isotype control antibody confirms the specificity of the primary antibody for its target protein.

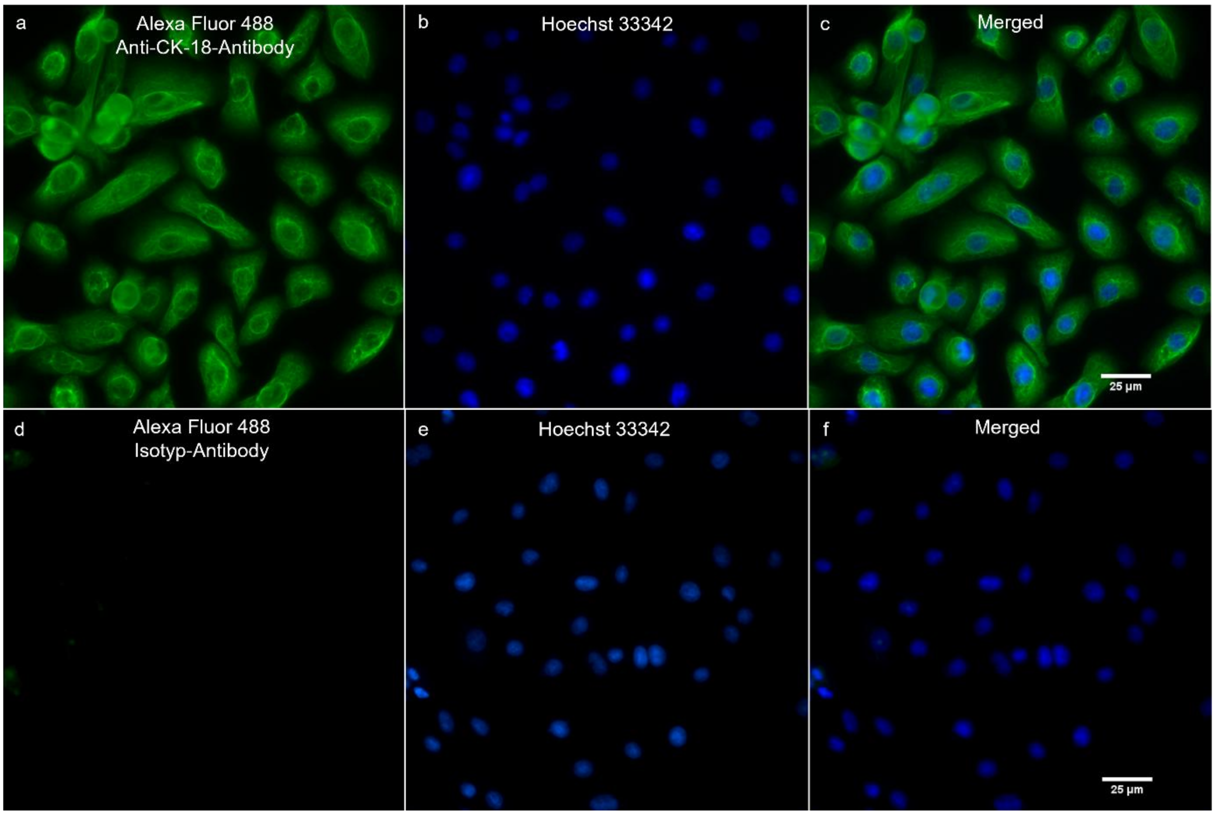


**Fig. S1** Representative epifluorescence images of hBECs following immunofluorescence with anti-CK18 antibody (**a-c**) or isotype control antibody (**d-f**), counterstained with Hoechst 33342. Images are shown as false-color representations (n = 6 experiments). Abbreviations: CK, cytokeratin.
